# Multi-attribute decision-making in macaques relies on direct attribute comparisons

**DOI:** 10.1101/2023.10.22.563329

**Authors:** Aster Q. Perkins, Zachary S. Gillis, Erin L. Rich

## Abstract

In value-based decisions, there are frequently multiple attributes, such as cost, quality, or quantity, that contribute to the overall goodness of an option. Since one option may not be better in all attributes at once, the decision process should include a means of weighing relevant attributes. Most decision-making models solve this problem by computing an integrated value, or utility, for each option from a weighted combination of attributes. However, behavioral anomalies in decision-making, such as context effects, indicate that other attribute-specific computations might be taking place. Here, we tested whether rhesus macaques show evidence of attribute-specific processing in a value-based decision-making task. Monkeys made a series of decisions involving choice options comprising a sweetness and probability attribute. Each attribute was represented by a separate bar with one of two mappings between bar size and the magnitude of the attribute (i.e., bigger=better or bigger=worse). We found that translating across different mappings produced selective impairments in decision-making. When like attributes differed, monkeys were prevented from easily making direct attribute comparisons, and choices were less accurate and preferences were more variable. This was not the case when mappings of unalike attributes within the same option were different. Likewise, gaze patterns favored transitions between like attributes over transitions between unalike attributes of the same option, so that like attributes were sampled sequentially to support within-attribute comparisons. Together, these data demonstrate that value-based decisions rely, at least in part, on directly comparing like attributes of multi-attribute options.

**Significance Statement:** Value-based decision-making is a cognitive function impacted by a number of clinical conditions, including substance use disorder and mood disorders. Understanding the neural mechanisms, including online processing steps involved in decision formation, will provide critical insights into decision-making deficits characteristic of human psychiatric disorders. Using rhesus monkeys as a model species capable of complex decision-making, this study shows that decisions involve a process of comparing like features, or attributes, of multi-attribute options. This is contrary to popular models of decision-making in which attributes are first combined into an overall value, or utility, to make a choice. Therefore, these results serve as an important foundation for establishing a more complete understanding of the neural mechanisms involved in forming complex decisions.

## Introduction

Complex, real-world decisions often do not have a clear best option. This is because there are typically multiple relevant features, or attributes, such as quantity, quality, or cost, that need to be considered in order to select the most preferred combination. This process could evolve in different ways, and it remains unclear how the brain uses information about different attributes to compute these types of complex decisions.

A canonical solution to this problem is to compute an integrated value, or utility, for each option from a weighted combination of all relevant information. This could allow different options to be compared on a common scale, accounting for context, internal state, and different features or attributes of unalike options (1–4). Many views have proposed that decision computation occurs by comparing these integrated option values (Figure 1A) (5, 6). Neural responses correlating with variables in these models have been widely reported (7–11), supporting the idea that the brain is capable of computing something like integrated value.

**Figure 1.**
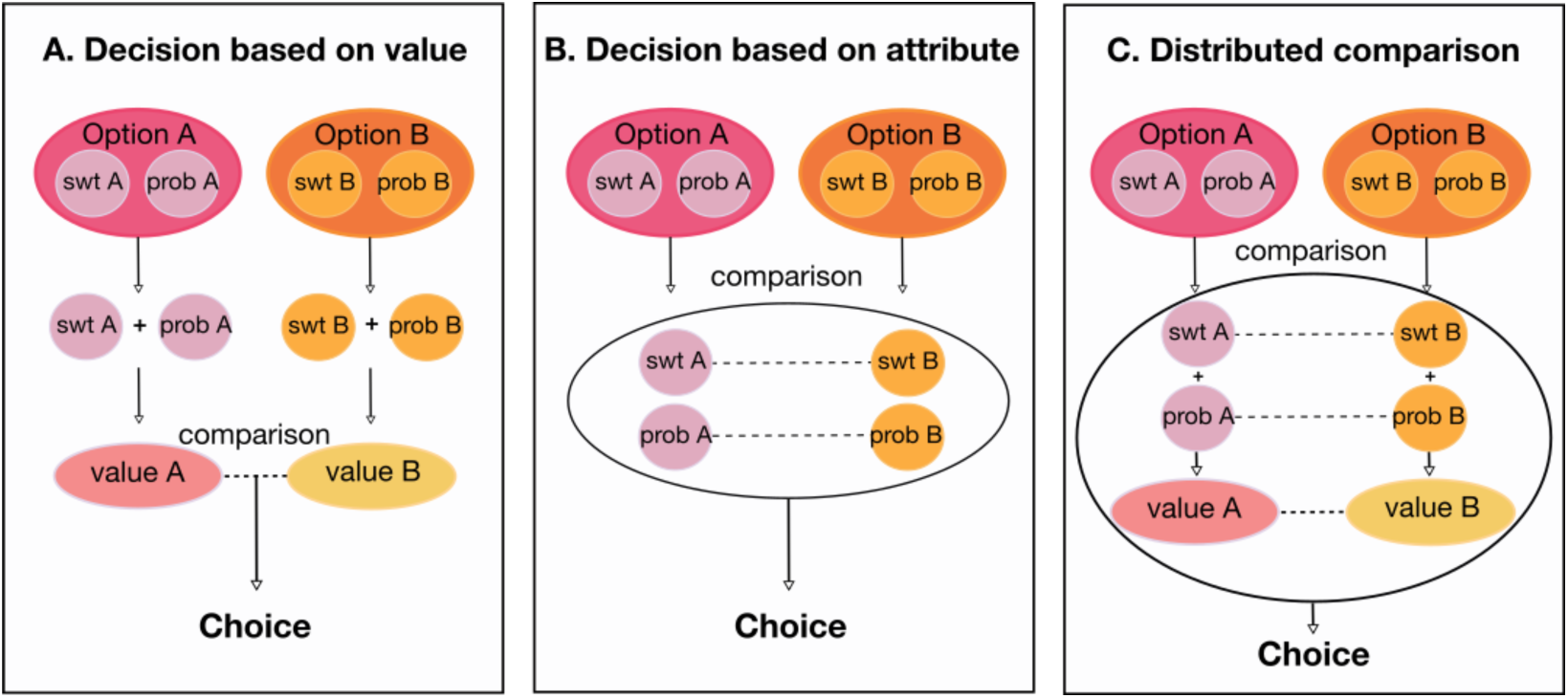
Three models of goal-directed decision-making. Each panel shows a representative two-option choice. As in our task, each option (A and B) consists of two attributes: sweetness (swt) and probability (prob). **A**. In the ‘integrate then compare’ model, attributes are combined to compute an integrated value for each option, which is then compared to the other option’s value to produce a decision. **B**. An attribute comparison model, in which like features (attributes) are compared directly, so that the number and weight of each competition in favor of an option produces a choice. **C**. A hybrid model, in which comparisons occur at the multiple levels, including comparisons of attributes and integrated values, so that decisions arise from distributed processes. (Figure adapted from (31)).

In contrast, behavioral anomalies that are inconsistent with the “integrate then compare” model of decision-making are also well documented (12–17) and suggest that integrated value may not be the sole input into the decision process. For instance, choice biases known as *decoy* or *context effects* arise when third options are added to the choice set (12–14). In these cases, the direction of the bias depends on where the third option’s attributes are situated with respect to those of the other options. Moreover, these effects can be explained by the dynamics of decision formation in models where comparisons take place at the level of individual attributes (Figure 1B) (15–17), (though see 18–22). These include decision field theory, which accounts for context effects by eschewing models that rely on overall utility, and instead suggesting that preferences evolve over the course of a decision by incorporating a continuous stream of comparisons of attributes (23, 24). This also accounts for the effects of attention, which influences both choice and neural encoding of values of visually attended items (25–27). Related views suggest that attribute processing may occur in parallel to the computation, and potentially comparison, of integrated values (Figure 1C). For instance, the distributed theory of decision-making posits that decisions are an emergent property of computations formed in multiple brain regions (28–30). Regions share information in an ordered way, but do not represent a circuit in the canonical sense. Rather, recurrent and overlapping computations that might be based on attributes or integrated values are spread across the brain and produce choice through their interactions. Collectively, these models represent alternatives to utility-based explanations of choice behavior and demonstrate that the mechanism of preference-based decision-making is far from a solved matter.

Here, we explored the potential for direct interaction or comparison between attributes by analyzing multi-attribute decision-making behavior in rhesus macaques. The monkeys were trained to choose between options that varied in the sweetness of a sugar water reward and likelihood of receiving it. Each attribute was represented by visually distinct bars whose sizes changed with respect to the sweetness and probability level. Critically, bars were presented in two mappings denoted by different colors: one in which the size changed proportional to the goodness of the attribute, and one in which the size changed inversely proportional to the goodness of the attribute. Both attribute magnitude and mapping varied independently, allowing us to reveal consistent patterns of choice deficits that indicated the spontaneous use of within attribute comparisons, even when all information was available to compute integrated values. Therefore, comparisons between individual attributes of multi-attribute options appear to be a natural feature of goal-directed decision-making in monkeys. This result opens new questions about the neural mechanisms underlying complex, goal-directed decisions.

## Results

### Monkeys use multiple attributes to make optimal choices

Two monkeys performed a decision-making task in which they selected between options comprising two attributes: the sucrose concentration of a fluid reward (*sweetness)*, and the probability that it would be delivered (*probability*) (**Figure 2A**). Attributes varied across 5 magnitudes, denoted by the length of separate bars. Bar size varied either directly (i.e., bigger is better) or indirectly (i.e., bigger is worse) with the magnitude of sweetness or probability, and the mapping was denoted by the bar color (**Figure 2B**). For example, a large blue bar would indicate a sweeter option, while a large pink bar would indicate a less sweet option. Sweetness and probability magnitudes and mappings varied independently in both options, and pairs of bars composing an option were pseudorandomly assigned to one of six possible locations on the task screen. To initiate a trial, subjects held a touch-sensitive bar while fixating a central point. Two options were shown, each consisting of two bars. Monkeys freely viewed the options, and made a choice by releasing the bar while holding gaze on one option. Reward was probabilistically delivered, depending on the option selected. We quantified choice behavior from 57,082 and 65,384 trials for Monkeys D and C respectively, performed over 69 and 76 sessions.

**Figure 2.**
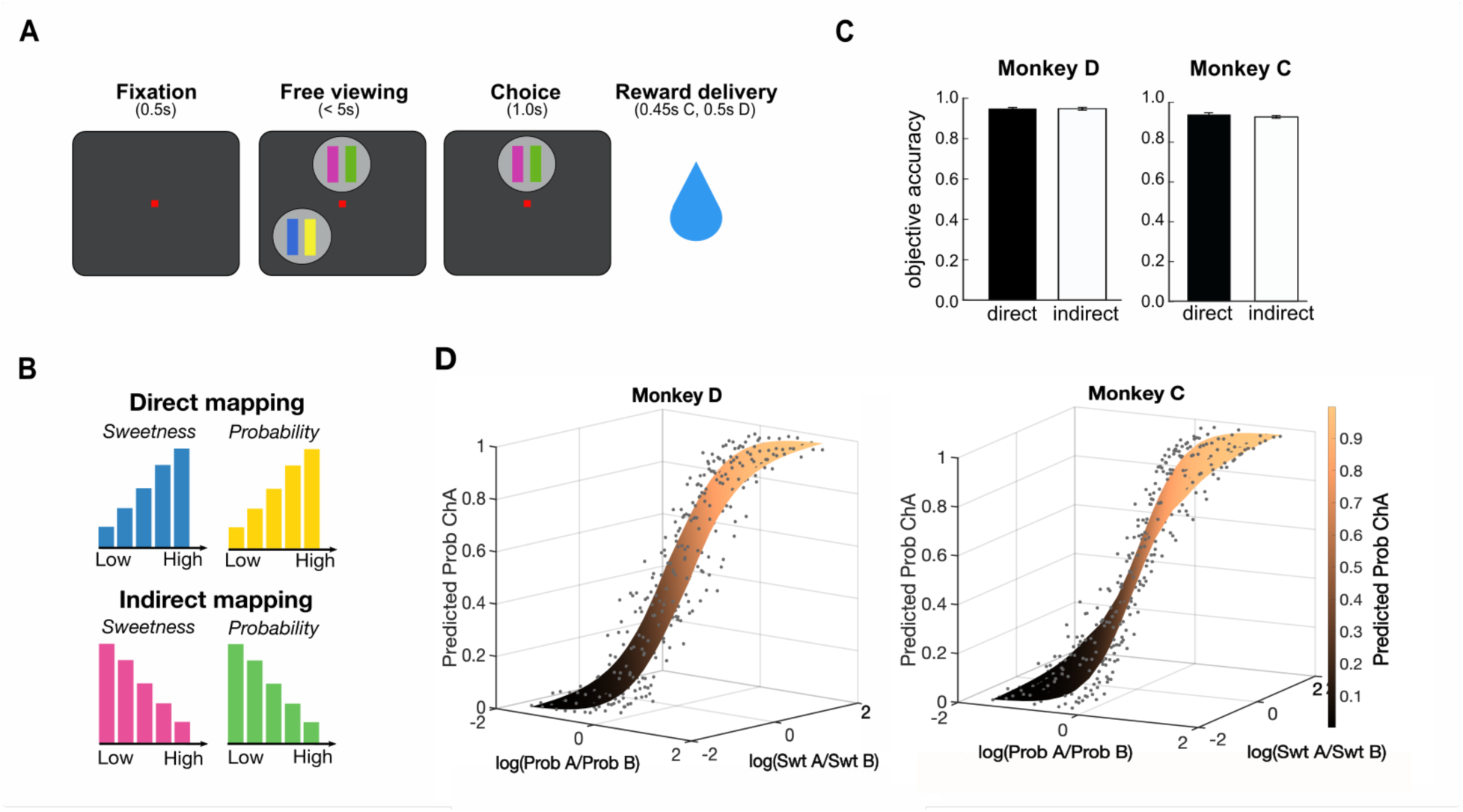
Multi-attribute choice task and performance. **A**. Trial schematic, where each option consists of a pair of bars indicating sweetness (left) and probability (right). The figure shows two of six possible option locations. After the monkey makes a selection, the unchosen option disappears and reward is probabilistically delivered. **B**. Each attribute bar had one of two mappings. Blue/yellow indicated a direct mapping, in which taller bars represented greater sweetness/probability. Magenta/green indicated an indirect mapping, in which taller bars represented lower sweetness/probability. **C**. Accuracy based on expected value in “objective trials,” split by direct and indirect mapping. Error bars = SEM. **D**. Interpolated predictions from fitted models (surfaces) show that the probability of choosing an arbitrary option A (Prob ChA) increases as the relative sweetness or probability of that option increases, and the attributes contribute roughly equally to decisions (Eqn. 3). Actual choice frequencies (points) are well fit by the model. D: N=57082, C: N=65384

Both monkeys performed the task well, and often chose the option with the highest expected value, when this was defined as the product of sweetness and probability (Across sessions: Monkey D: 0.81 ± 0.007 (95% CI); Monkey C: 0.82 ± 0.006 (95% CI)). Subjects chose the higher value option slightly less frequently on trials where all attribute mappings were indirect, compared to when they were all direct (Across sessions: Monkey D: direct: 0.84 ± 0.013 (95% CI), indirect: 0.81 ± 0.02 (95% CI). Monkey C: direct: 0.84 ± 0.013 (95% CI), indirect: 0.80 ± 0.013 (95% CI)). However, this effect was small and monkeys still performed at a high level for both mappings, demonstrating that they are capable of using both mappings flexibly to retrieve information about the different attributes.

Because this definition of expected value does not account for subjective preferences that differentially weight sweetness or probability, we also calculated objective accuracy on trials where one option was superior in both sweetness and probability. This excluded trials in which one option was greater in one attribute but lower in the other (note that this is irrespective of mappings). Objective accuracies were very high, likely reflecting the fact that these were easier decisions (**Figure 2C**. Across sessions: Monkey D: 0.92 ± 0.008 (95% CI), Monkey C: 0.93 ± 0.013 (95% CI)). The effect of direct/indirect mapping on accuracy also vanished in the objective trials for Monkey D, and was mitigated for Monkey C (Across sessions: Monkey D: direct: 0.95 ± 0.012 (95% CI), indirect: 0.95 ± 0.014 (95% CI), Monkey C: direct: 0.94 ± 0.02 (95% CI), indirect: 0.92 ± 0.013 (95% CI)).

To quantify the subjects’ preference weightings for the two attributes, we fit choice data with a multinomial logistic regression similar to those used in previous decision-making tasks (32, 33). The model predicted choices from the log-ratio of the ordinal magnitudes of each attribute, the mapping of each attribute (direct/indirect), and the X-Y positions of each option on the task screen (see Methods). The magnitudes of both attributes strongly predicted choices in both subjects (**Figure 2D**), showing that monkeys tended to pick the option with higher sweetness and probability. They also weighted the attributes roughly equally, with a slight bias toward probability (Monkey D: Log(ProbA/ProbB) β=1.83, Log(SwtA/SwtB) β=1.62. Monkey C: Log(ProbA/ProbB), β=2.64, Log(SwtA/SwtB) β=1.18. All p<1×10^-13^) (**Supplemental Figure 1 & 2A**). In contrast, attribute mappings and location on the screen had minimal effect on choice. Monkey D showed a tendency to select options with indirect probability mappings in early testing sessions, but this disappeared over time (**Supplemental Figure 2B)**. Therefore, monkeys primarily used sweetness and probability information to guide decisions.

### Mapping mismatches impair attribute comparisons

Next, we considered two operations that could be involved in computing decisions: integrating attribute values within an option, and directly comparing like attributes across the different options. Because it is relatively straight-forward to identify a larger or smaller bar from a pair, these processes should be easier when bars have the same mapping, and more difficult when they differ (i.e., when one is direct and one is indirect). If options are assessed solely on the basis of integrated value, the attributes need to be integrated into an overall value prior to comparison with another option. In this case, mismatching direct and indirect mappings *within an option* should disrupt the integration of the attributes and impair choice accuracy. Once the option values are computed, the original bar mapping should no longer matter, meaning that mismatches across options should not have the same effect. Conversely, if monkeys use direct attribute comparisons to compute a choice, then mismatched mappings *within like attributes* (across options) should impair choice accuracy (**Figure 3A**). Because attribute mappings varied independently, we were able to sub-select trials where attributes were either matched or mismatched in different combinations to test these hypotheses.

**Figure 3.**
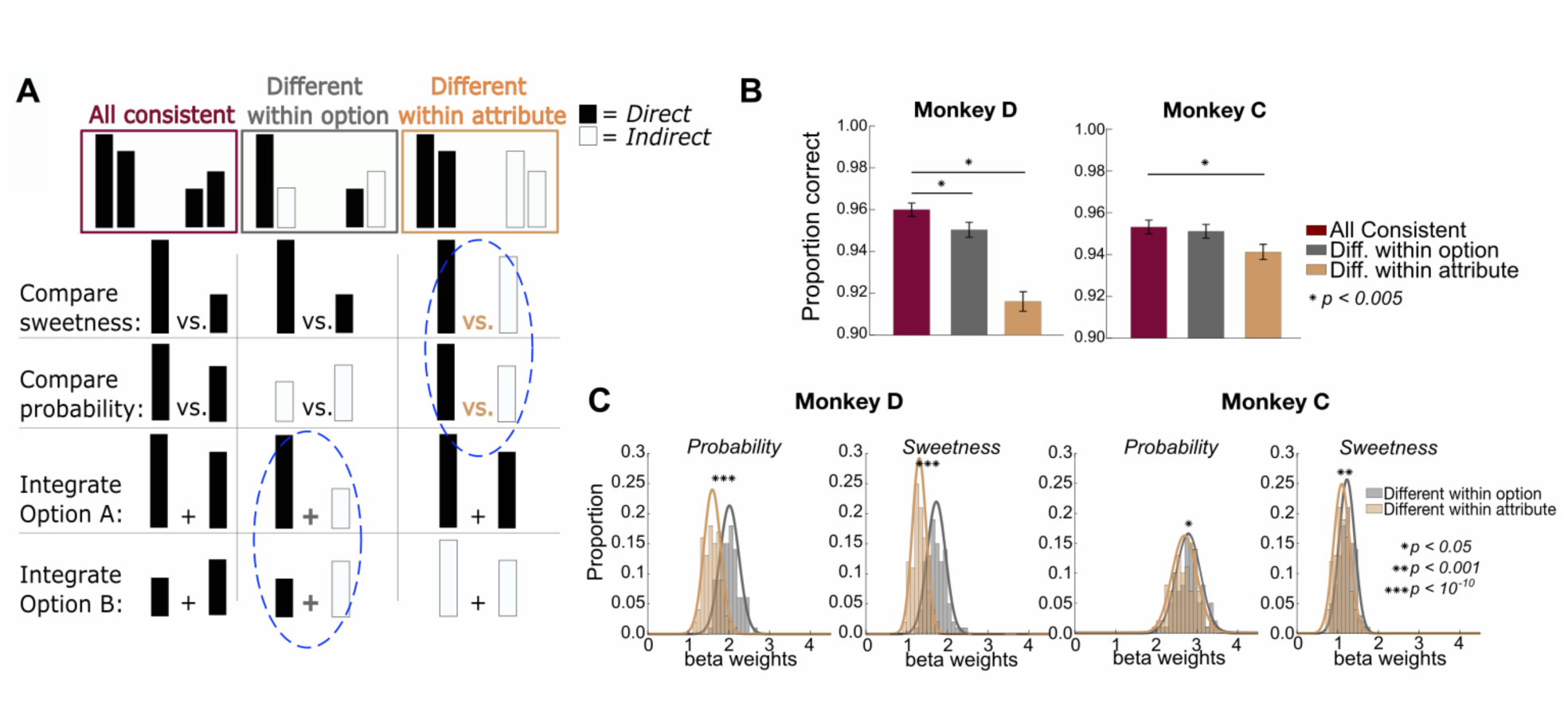
Choice accuracy varies with attribute arrangement. **A**. Schematic of three trial types. Burgundy: all attributes have the same mapping, either all “direct” (shown) or all “indirect” (not shown). On these trials, comparison or integration processes would take place among attributes with the same mapping. Gray: mappings differ among attributes within an option, but like attributes of the two options are the same. In this case, attribute comparisons would take place between bars with the same mapping, but integration within an option would take place across mismatched mappings (blue dashed circle). Gold: mappings differ within like attributes but are the same within each option, so that attribute comparisons would take place between mismatched mappings (blue dashed circle), but value integration would take place between bars with the same mapping. Note that in the last two trial types, there are always two bars of each mapping, and only their arrangement differs. **B**. Accuracy on trials of each type in which there was also an objectively better option (n = 3728 and 4188 trials for Monkey D and C respectively). Accuracy was consistently lower when like-attributes differed (gold). One-way ANOVA: Monkey D: F_2,10897_=35.58, p=3.97×10^-16, Monkey C: F_2,12595_=3.48, p=0.0308. * = significant post-hoc comparisons. Error bars = SEM. **C**. Distributions of bootstrapped samples of 400 trials, sampled from all trials (not just those with an objectively better option) in which attribute mappings were different within option (gray), or different within attribute (gold) as in **A**. Mismatched mappings within attribute consistently resulted in smaller slopes (i.e., more variable choices). * significant Wilcoxon rank-sum tests. D: N=57082, C: N=65384

First, we focused on trials with an objectively better option, as described above, so we could calculate choice accuracies. Among these trials, we compared accuracy when all four bars had a consistent mapping, meaning all were direct or all were indirect, to accuracies when two bars were direct and two were indirect. In the latter case, we separated trials in which like attributes had the same mapping but differed within an option, and those in which the two bars of an option had the same mapping but differed from the two bars of the other option. On these trials, monkeys were consistently less accurate when bars of like attributes had mismatched mappings, suggesting a disruption of attribute-level comparisons (One-way ANOVA: Monkey D: F_2,10897_=35.58, p=3.97×10^-16^, Monkey C: F_2,12595_=3.48, p=0.03) (**Figure 3B**). In contrast, when options are matched within attribute, but mismatched within option, we found less or no deficit in choice accuracy. Because there were the same number of direct or indirect attributes in each condition and attributes merely varied in arrangement, the slightly lower accuracy using indirect bars cannot explain this result. Likewise, mappings were randomly and independently assigned to each attribute and option, so that the mapping of a particular attribute could not influence this effect. Instead, the arrangement of attribute mappings in the choice determined choice accuracy. Similar but more subtle effects were found in reaction times (**Supplementary Figure 4**).

To assess effects across all choices (not just objective ones), we separately modeled trials in which bar mappings differed within an option and those in which bars differed within like attributes as sigmoid functions (***Eqn 3***). If different bar mappings impeded within-attribute comparisons, we would expect less consistent choices on those trials, quantified as shallower fitted slopes. To control for effects of trial number, we created 100 bootstrapped samples of 400 trials for each condition and estimated distributions of fitted slopes for each. We found that choices were more variable when bars differed within attribute versus within option for both sweetness and probability (**Figure 3C**). Therefore, preventing monkeys from easily making direct attribute comparisons resulted in less accurate choice behavior, suggesting that they rely on these comparisons to make multi-attribute decisions.

### Gaze patterns during multi-attribute choices

As humans or monkeys decide, they acquire information by shifting gaze among visible options, and this process of allotting attention influences the dynamics of decision formation (26, 34, 35). Our task presented attributes as physically separate bars, which allowed us to use eye tracking to quantify gaze patterns across the different options and attributes. First, we focused on fixations on attribute bars before a choice was made (the pre-choice epoch). The final fixation that coincided with the choice report, when the monkey released the touch bar, was excluded. Monkey C made slightly fewer fixations per trial than Monkey D (D: mean = 2.27 fixations, SE = 0.0059; C: mean = 1.67 fixations, SE = 0.0035), and both monkeys exhibited idiosyncratic preferences for looking at either the sweetness (Monkey D) or probability (Monkey C) bars (**Supplemental Figure 5**). Notably, despite these tendencies, behavioral choices reflected clear usage of information represented by both attribute bars (see **Fig 2D**). Therefore, gaze bias is not a one-to-one proxy for preference and is dissociable from choice behavior, although they are not entirely independent.

First, we determined whether there were general patterns that related gaze behavior to choices. Both monkeys made more fixations per choice as the difference between the attributes in the two options decreased, so that they looked more at the options when choices were more difficult (i.e., close in value, **Figure 4A**). We quantified this with a general linear regression that predicted the number of fixations on any attribute bar from the difference in offered sweetnesses and probabilities, as well as the mapping of each attribute, and found that smaller differences in either attribute predicted more fixations per trial (Monkey D, C: sweetness difference: *β*=-0.11, −0.02, *p*<10^-99^, 10^-6^; probability difference: *β* =0.17, −0.11, *p*<10^-263^, 10^-244^). In addition, consistent with previous reports (25), fixations were longer when monkeys looked at either bar of the option they would eventually select (the *chosen option*), compared to the one they would not (the *unchosen option*) (**Figure 4B**). Therefore, aspects of the monkeys’ gaze patterns aligned with expected effects of value and choice.

**Figure 4.**
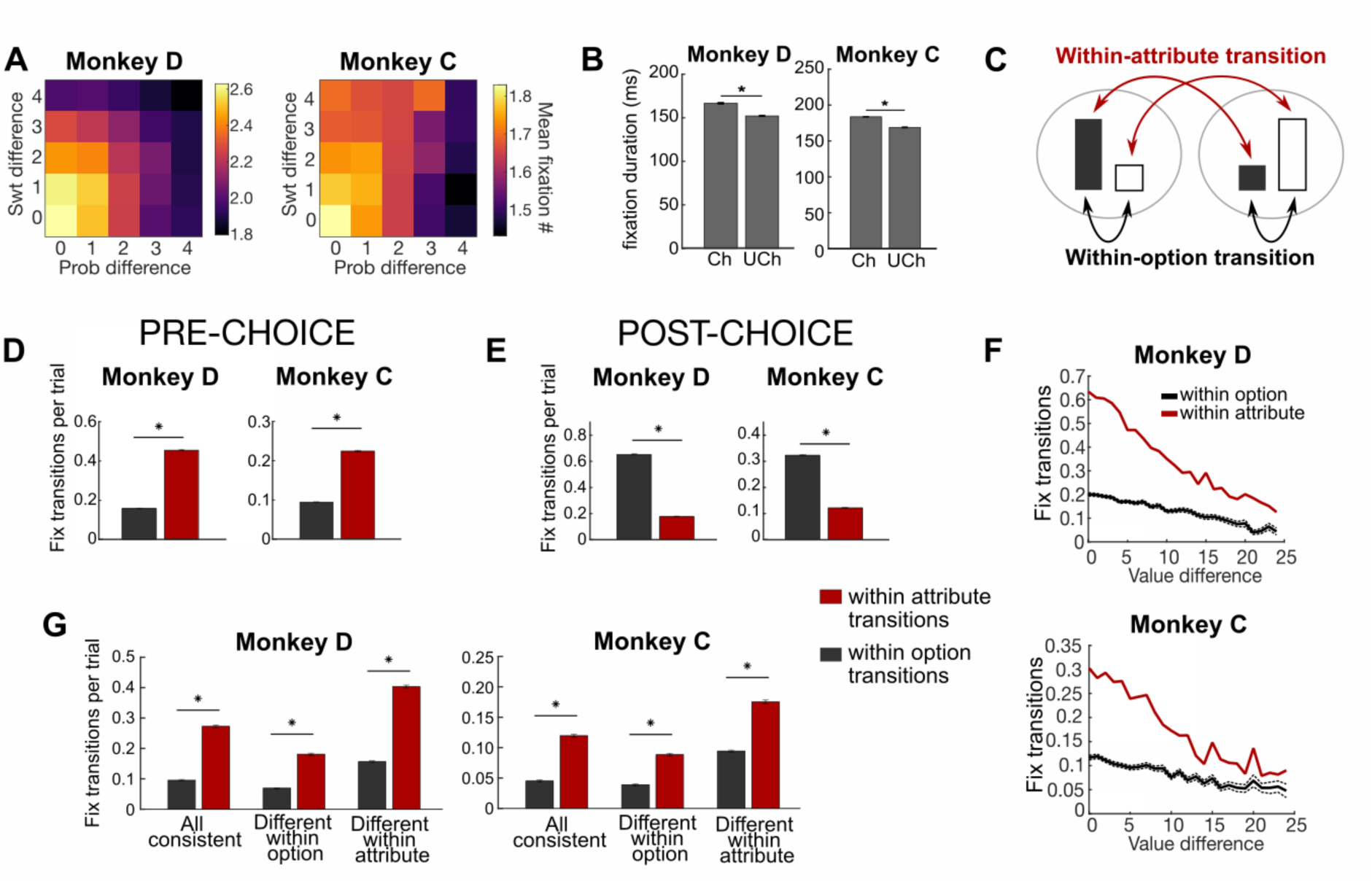
Gaze patterns in multi-attribute choices. **A**. Colormaps show that the mean number of fixations on any bar scaled with the ordinal difference between either sweetness or probability of the choice options. D: N= 51115, C: N= 53727. **B**. The mean duration of fixations on either bar of the chosen option (Ch) was higher than those on bars of the unchosen (Uch) option. Bars show the aggregate of all fixations in a trial, and error bars represent 99% confidence intervals. D: N= 51115, C: N=53727. **C**. Schematic showing within attribute (red arrows) and within option (black arrows) gaze transitions. Black bars represent sweetness, white bars represent probability, and an option comprises two bars in the same gray circle. **D**. Number of fixation transitions within-option (black) or within-attribute (red) for each monkey before a choice is made. Error bars = SEM. * p < 10^-10^. N=88819 (D) and 106800 (C) (fixation transitions). **E**. Number fixation transitions as in **D**, except after the choice is reported. N=81909 (D) and 110400 (C) (fixation transitions). **F**. The number of fixation transitions per trial within-option or within-attribute varied with the difference in expected value of the choice options on that trial. Dashed lines show SEM (very small due to high number of trials). N=43430 (D) and 27055 (C) (trials). **G**. The number of fixation transitions per trial, split by attribute arrangement, as in Figure 2E. Planned post-hoc comparisons from two-way ANOVA compared within attribute and within option transitions. * p < 0.0033 (Bonferroni corrected). Error bars = 95% CI. N= 43430 (D) and 27055 (C) (trials).

Next, we assessed effects of attribute mapping on gaze patterns. Here, Monkey D slightly preferred to look at indirect sweetness and probability bars, while Monkey C preferred to look at indirect sweetness bars, but direct probability bars (**Supplemental Figure 5**). These patterns are reflected in the effects of bar mappings on fixation number in the linear model (Monkey D, C: number of indirect swt bars: *β* =-0.02, 0.02, *p*=0.03, *p*<10^-6^; number of indirect prob bars: *β* =-0.04, −0.08, *p*<10^-5^, 10^-57^), but effects are weak compared to the attribute magnitudes and idiosyncratic to the individual monkeys.

Finally, we assessed the same effects on fixation durations, using a linear regression model to predict the duration of any fixation from features of the fixated bar (magnitude and mapping), features of the non-fixated bars, and fixation sequence. Across monkeys, fixations were longer for probability bars, chosen options, indirect bars, and greater values of the fixated attribute (**Supplemental Figure 6**). In addition, there were some effects of unfixated attributes. Both monkeys showed shorter fixations when either attribute of the other option had a greater value. However, for greater values of the other attribute of the same option, Monkey C had longer fixations while Monkey D had shorter. Together, the strongest patterns in gaze behavior were related to the monkeys’ choices, with only weak or idiosyncratic effects of other variables such as attribute mapping, suggesting that gaze can provide a window into decision-making processes.

### Gaze transitions are consistent with attribute comparison

Our choice data suggested that monkeys compared like attributes of the two options when making a decision. Gaze shifts can provide insights into the order of information sampling, and therefore reveal patterns consistent with attribute comparison or value integration. If monkeys use direct comparison of attributes to help them decide, we predicted that most gaze transitions would be between bars representing like attributes. To assess this, we determined how often gaze transitioned between attributes of the same option (i.e., sweetness bar A Η probability bar A or sweetness bar B Η probability bar B), which would be expected during integration into an overall value. We compared this to transitions between two bars representing like attributes, (i.e., sweetness bar A Η sweetness bar B, or probability bar A Η probability bar B), which would be expected with attribute comparisons (**Figure 4C**). For this analysis, trials with less than two fixations before a choice were excluded, yielding 43430 (D) and 27055 (C) trials for gaze transition analyses.

Both subjects made significantly more within-attribute transitions than within-option transitions in the pre-choice epoch (Monkey D: t_88819_ = −83.27, *p*<1×10^-13^. Monkey C: t_106800_ = −57.77, *p*<1×10^-12^) (**Figure 4D**). This is consistent with the idea that monkeys compare information about the like attributes to each other. As a follow-up, we quantified the same measure in the post-choice epoch, when only the chosen option was displayed on the screen. Despite the unchosen option not being visible, monkeys occasionally looked to the location where it previously was, but predominantly we found within-option transitions as expected (Monkey D: t_81909_ = 113.60, *p*<1×10^-13^, Monkey C: t_110400_ = 77.56, *p*<1×10^-13^) (**Figure 4E**). In summary, monkeys showed within-attribute sampling of information prior to making a choice.

Next, we assessed how both types of gaze transitions changed with choice difficulty, using the difference in expected values between the two options as a proxy for the difficulty of the decision. While both within-option and within-attribute transitions increased on more difficult trials, the increase in within-attribute transitions was greater than within-option transitions (**Fig. 4F)**. This was quantified by significantly greater slopes in a regression that correlated number of within-attribute gaze transitions and expected value difference, compared to the same regression for within-option transitions (t-test for differences between slopes. Monkey D: t_113160df_=-29.51, *p*<1×10^-11^, Monkey C: t_126448_=-20.89, *p*<1×10^-10^). Therefore, within-attribute transitions predominate when choices are more difficult, which would be expected if these shifts in attention mediate comparison of like attributes.

Finally, we considered whether gaze transitions also reflected the perceptual difficulty of the decision by assessing trials where the bar mappings were 1) all consistent, 2) different within option, or 3) different within attribute, as in **Figure 3A**. Here, monkeys primarily made within-attribute gaze transitions, regardless of attribute arrangement. There was a significant interaction between categorical factors of trial type (attribute mapping: all consistent, different within option, or different within attribute) and transition type (within-attribute or within-option) on the number of fixation transitions per trial (Two-way ANOVA. Monkey D: F_2,21536_=33.32, *p*=3.58 x 10^-15^, Monkey C: F_2, 23944_=5.02, *p*=0.0066). This was driven by within-option transitions increasing proportionally more on different within attribute trials (i.e., when mappings were consistent within option), although there were more within-attribute transitions on all trial types (**Figure 4G**). Therefore, in agreement with patterns in choice behavior, gaze behavior also indicated that monkeys used attribute comparisons to guide their choices. In addition, gaze patterns primarily reflected choice difficulty related to value difference but not perceptual difficulty.

## Discussion

A central premise of most decision-making models is the idea that choices involve the computation and comparison of integrated values. However, other views have proposed that unintegrated attributes also play a role. Here, we used a novel multi-attribute decision-making task for monkeys to test whether complex decisions incorporate information about unique attributes as part of the choice process. Patterns in choice accuracy, preference consistency, and directed gaze all indicated that monkeys spontaneously used direct comparisons of like attributes to arrive at a decision. Importantly, this occurred while information was also available to compute integrated values, so that the task revealed aspects of the monkeys’ natural tendency to use unintegrated attributes. Although these data do not exclude a role for integrated value comparisons, they do emphasize that integrated value is not the sole decision variable that the brain uses when computing a preference-based choice.

Our results provide support for a class of model in which choices are based on evaluation of individual attributes, either without (24, 36–38) or in parallel to (28, 29) the computation of integrated values. Many models that include within-attribute comparisons also account for classic anomalies in multi-attribute choices, such as context effects (39, 40). For example, one type of model proposes that attributes are compared in a pairwise manner and evidence in favor of an option accumulates by linearly combining these with attentional weights. This accounts for attraction effects because greater attentional weights are placed on options that are closer together in attribute space (36, 39). In addition to our results, models such as this support the validity of attribute-based theories of decision-making.

Our results could also align with theories that propose decisions arise from parallel comparisons of multiple variables, including like attributes, integrated values, and attribute saliencies (28, 30). From this view, decisions arise from distributed brain systems, which may be arranged hierarchically, such that information about option and attribute values can be jointly used along with other variables to produce a decision (28). Notably, our study cannot address situations where options do not share a common attribute. While some have proposed that this is a context in which integrated value comparisons are particularly important (2), other models solve this dilemma with a series of accept/reject decisions at the level of attributes (16). Regardless, our results demonstrate the need to consider attribute-level processes as a mechanism involved in multi-attribute decision-making.

Our conclusions are based in part on subtle impairments in choice behavior revealed by our task design, which incorporated different perceptual mappings of visual cues to attribute values (i.e., ‘bigger is better’ versus ‘smaller is better’). When mappings of two attribute bars were the same, the monkeys could compare them with a simple perceptual judgement to determine which is taller or shorter, or they could reference internal representations of either the specific meaning (e.g., 50% probability of reward) or the position on a value scale indicated by each bar. When the mappings differed, however, they couldn’t use simple perceptual judgements and had to rely on and translate between internal representations, making the mismatched condition potentially more difficult. Indeed, we found lower objective accuracies and less consistent preferences when mappings were different within like attributes, suggesting that like attributes were the components of the display that the monkeys were translating between.

Having the same mapping not only provides an easier way of comparing bars, but can also be seen as grouping them, either perceptually (by color) or informationally (by relationship of size to goodness). Previous work has shown that grouping attributes by type or the option they belong to, either through colored backgrounds or proximity, without introducing significant processing costs, does not change decision-making strategies (41). This makes it unlikely that the perceptual grouping drove our behavioral effects. Rather, our data suggest that an inefficiency was introduced to the decision process when the subjects had to compare attributes across informational groups (i.e., different mappings). Because the same degree of deficit was not seen when there were discordant informational groups within an option, it suggests that operations taking place across this grouping are less critical to the choice process, and the brain relies more on direct attribute comparison to make decisions.

It is unlikely that our behavioral effects were driven by perceptual features in other ways, for instance by way of differences in preference or accuracy of value estimation for one of the mappings. This is because we subselected trials from our large data set that had an identical number of “direct” and “indirect” bars on the screen in each condition (i.e., different within option or different within attribute). With this approach, only the particular arrangement of the matched or mismatched attributes varied across conditions, eliminating effects that might arise merely from the monkeys’ interpretations of the direct or indirect condition.

If monkeys used direct comparison of attributes to help them make decisions, we expected this would also be evident in the way they gathered information while making a choice. For this, we analyzed patterns of fixation, and predicted that gaze would predominantly transition between the bars representing like attributes. From this view, shifting gaze reflects a process by which items are brought into the focus of attention, and therefore sequentially sampled (25). As predicted, we found a tendency to shift gaze between like attributes, particularly on more difficult choices when the options were more similar in value. This is consistent with human studies, which find that gaze tends to shift primarily between alternatives along single attribute dimensions during multi-attribute choices (42–44). It is also in-line with lesion evidence indicating the importance of sampling attributes in sequence in choice behavior. For instance, when asked to choose between hypothetical apartments that vary in multiple features, control subjects tended to favor attribute-based strategies, and uncovered information about like attributes in sequence (45). In contrast, individuals with damage to the ventromedial prefrontal cortex (vmPFC), an area important for value-based decision-making, tended to favor alternative-based decision strategies, seeking out all of the information pertaining to a given option at once. This suggests that damage to decision-making circuits disrupted an otherwise common tendency to sequentially gather information about like attributes. Therefore, across species, gaze patterns consistently reveal preferences for within-attribute gaze transitions, reflecting a search strategy oriented toward making a within-attribute comparison.

Because the same decision can be made a number of different ways, and can rely to varying degrees on unobservable internal states, studies often seek a fuller understanding of decision processes by assessing the neural substrates (46). A number of brain regions have been implicated in value-based decision making. Among these, the orbitofrontal cortex (OFC) is particularly important, as damage or disruption consistently alters value-based choice behavior, suggesting that OFC neurons perform choice-relevant computations (47, 48). Integrated value signals are commonly found within OFC, including in single unit firing rates (7–9), population codes (49, 50), field potentials (50–52), and fMRI BOLD signals (53, 54), and this has been taken as evidence that integrated value is the key decision variable in OFC. However, multiple labs consistently report neurons in monkey OFC (primarily area 13) that encode the value of unique attributes (7, 55–59), and similar signals can be found in human fMRI BOLD (60). These responses are understudied compared to the more prevalent integrated value signals, making it unclear whether they inform choice behavior or are used to compute the integrated values that then guide choice. Based on our results, the possibility of attribute-level processing should be considered when interpreting neural correlates of choice behavior in OFC and other regions.

In summary, our results provide robust behavioral evidence that monkeys use attribute comparisons when making multi-attribute decisions. This stands in contrast to models in which choices emerge exclusively from the computation and comparison of integrated values, and instead supports those that rely at least in part on attribute-level operations. This is particularly relevant when interpreting neural responses in choice paradigms where decision variables, such as attribute values and integrated values, are often correlated. Without directly testing alternate possibilities, the same data could be interpreted as support for entirely different psychological and neural mechanisms of decision-making.

## Materials and Methods

### Subjects and behavior

Two adult male rhesus macaques (“C” and “D”) participated in the experiment. Neither animal had been used for previous studies; however, they had been trained on a battery of tasks. They were each 5 years old and weighed 11.8 kg and 9.4 kg respectively at the start of experiments. Each was surgically implanted with a titanium head positioner. All procedures were in accord with the National Institutes of Health guidelines and were approved by the Icahn School of Medicine at Mount Sinai Animal Care and Use Committee.

During experiments, monkeys sat in a primate chair in a darkened testing chamber, and were head-fixed facing an 18-inch computer monitor positioned 17 inches away from the subjects’ faces. MonkeyLogic software (61–63) controlled the behavior interface. Subjects’ eye position was continuously monitored by an infrared eye tracker (ViewPoint Eyetracker USB-400, Arrington Systems) at a sampling rate of 400Hz. This signal was acquired by MonkeyLogic at 500Hz to align to behavioral data.

### Task

The monkeys were trained to perform a multi-attribute decision-making task. Each trial began with the appearance of a fixation cue in the center of the screen. When the monkey gazed at the fixation point and simultaneously held a touch-sensitive bar for 550 ms, two (80% of trials) or three (20% of trials) choice options were displayed on the screen. Options were shown at 2 or 3 of 6 potential positions in a hexagonal arrangement around central fixation. Option positions were selected randomly, with the constraint that they were never in adjacent positions on the hexagon. Only 2-option trials were analyzed in the present study.

Options were presented as a set of two bars. The width of each bar was 2 degrees of visual angle, and the height varied from 2 to 10 degrees. The size of the left bar of each pair indicated the sweetness level (25, 50, 75, 100, or 125 mM sucrose solution) while the right bar represented the probability level (30, 40, 50, 60, or 70%). The colors of the bars indicated whether the size of the bar increased with increasing reward/probability level (direct mapping), or decreased with increasing reward/probability level (indirect mapping). For example, a blue sweetness bar indicated a direct correlation (large bar = high sweetness), while a pink sweetness bar indicated an indirect correlation (large bar = low sweetness). Bar size and mapping varied randomly and independently on each trial.

While the monkey continued to hold the touch bar, he had up to 5 seconds to freely view the images while gaze position was tracked. They could make a selection at any time by holding gaze on any part of the desired option and releasing the touch sensitive bar. This would trigger the reward delivery immediately after an option was selected. If an option was not selected in this window, or if the touch bar was released when gaze was not directed toward an option, this triggered a 5s timeout, during which no reward was delivered and the screen displayed a red background. When this timeout was complete, the next trial could be initiated. Inter-trial intervals were 1s.

Rewards consisted of 0.33mL of fluid delivered over 500ms for Monkey D, and 0.297mL delivered over 450ms for Monkey C. Different amounts were titrated during pretraining to account for each subject’s relative weighting of sweetness and probability, in order to balance as closely as possible their subjective preference between the two attributes.

#### Quantifications and statistical analyses

The number of subjects was not predetermined by any statistical methods. Two animals is a common standard for monkey experiments because it is the smallest number with which we can demonstrate reproducibility. All of our analyses were carried out within subject, so that the *n* for statistical tests is number of trials or sessions. The same analyses were then carried out as replications in the second animal.

Some trials were excluded because the animal failed to make a choice within 5s (Monkey D: 599, 1.04%, Monkey C: 1203, 1.8%). There were more omissions on trials with a smaller difference in values (i.e., more difficult trials). Additionally, 3-option trials were part of the task design, but were excluded for the analyses in this paper. All statistical analyses were conducted with custom MATLAB (Mathworks) scripts.

##### Accuracies and reaction times

Accuracies were calculated as the number of trials in which the option with the highest expected value (defined as ordinal reward level multiplied by ordinal probability level) was selected over other options, divided by the number of valid trials in the same category. Because this calculation relies on assumptions about how the attributes are weighed by the subjects, we also computed accuracies for a subset of “objective” trials, in which one option had higher values of both sweetness and probability than the other (e.g., SwtA>SwtB and ProbA>ProbB), meaning it was the superior option regardless of attribute weighting. Reaction times were measured as the time from stimulus onset to release of the touch bar, indicating option selection. For Figure 3B, two 2-tailed binomial tests were performed between the “all consistent” group and the “different within option” group, and between the “all consistent” group and the “different within attribute” group (64). Significance levels were Bonferroni corrected for multiple comparisons to α = 0.005.

##### Choice regressions

We used the choice behavior of each monkey to examine the influence of task variables on decision-making. To quantify how each monkey weighs sweetness and probability, choices were modeled as a function of the log-ratio of each attribute (32, 33):

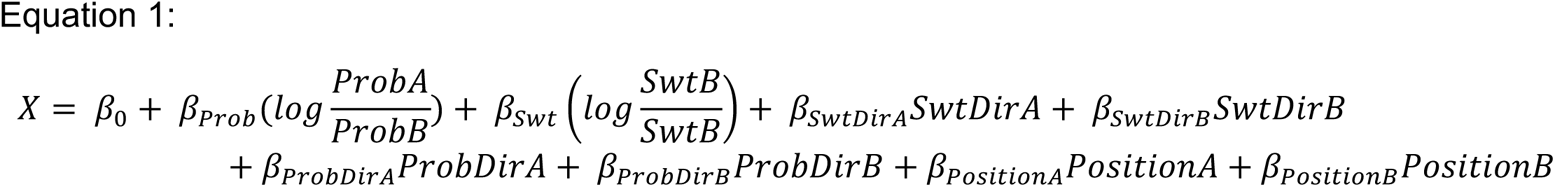

Options were arbitrarily designated A and B. *ProbA* or *B* and *SwtA* or *B* are the ordinal magnitudes of probability and sweetness available in each option. β_Prob/Swt_ are fitted weights of probability/sweetness attributes. β_0_ is a constant to capture any bias toward A or B. The probability of choosing option A was then calculated by a sigmoid function (*Eq. 2*).

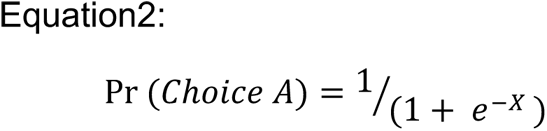

Choice probability colormaps were created from interpolating choice probabilities from the multinomial logistic regression on choice data, as above. Regressions were performed on concatenated data from all sessions, although session-by-session performance on each regression was assessed and is noted where included.

To perform the bootstrap analysis in Figure 3C, we reduced the Equation 1 to just the Sweetness and Probability ratios, as there were limited effects of the additional predictors in the original model:

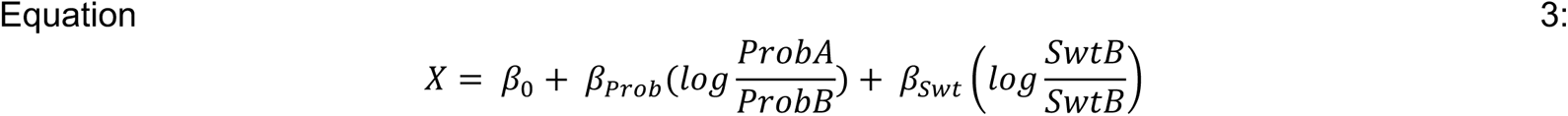

Eqn. 3 was then used with Eqn. 2 to perform the bootstrapped logistic regression.

##### Reaction time regression

To quantify how task variables affected reaction times, a general linear regression was performed on concatenated data from all sessions.

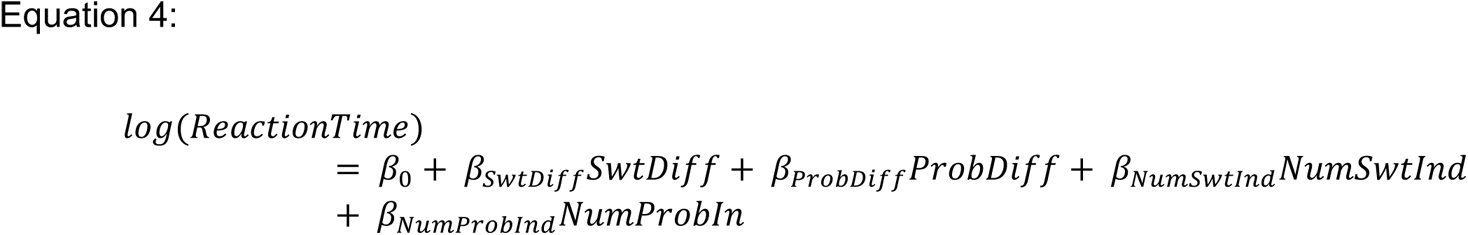

SwtDiff denotes the difference in the ordinal sweetnesses of the two options (0–4), and ProbDiff denotes the difference in the ordinal probabilities of the two options (0–4). NumSwtInd denotes the number of sweetness bars with the indirect mapping (0–2); NumProbInd denotes the number of probability bars with the indirect mapping (0–2). All predictors were mean centered (ranges above are pre-mean-centering).

##### Gaze data analysis

Gaze analyses were done with the EyeMMV Matlab package (65), which extracts fixations from continuous eye movements. Minimum fixation duration was set as 50 ms, a liberal criterion that allowed us to capture shorter fixations as assessed by visual inspection (66). Tolerances were set as t1=2 and t2=1, where t1 was the criterion in Euclidean distance within which gaze track records would be included in a fixation cluster, and t2 was the criterion in Euclidean distance within which gaze track records would be included in the computation of a fixation cluster mean. Regions of interest were then defined to include each attribute bar in a session, and fixations falling on each attribute were retained for analysis. The eye tracker was calibrated at the start of each session, and gaze data were further aligned in post-processing by centering data to the median x-y coordinates for the initial fixation window (placing it at the center fixation cue) across the session. Additionally, regions of interest around each attribute bar used for fixation detection were expanded by 0.75° of visual angle on the outside of the bars, and 0.5° of visual angle on the inside (so that there was no unassigned space between the bars), to capture fixations that were on the bar edges. Likewise, 0.5° of visual angle were added to the top and bottom of the regions of interest.

Unless otherwise stated, analyses were performed on fixations that occurred between option presentation and choice. Fixations detected immediately following option presentation were still at the center fixation cue, and were eliminated by removing all fixations beginning in a 50ms window following the stimulus presentation. The final fixation in the choice window was the fixation of the chosen option, and was also eliminated by removing the last fixation in the response window. Ultimately, 66 trials in Monkey D (0.09%) and 55 trials in Monkey C (0.065%) were excluded for having no detected pre-choice fixations.

##### Regressions on number of fixations

To quantify the effects of task variables on the number of fixations in each trial, we made a linear regression model as follows:

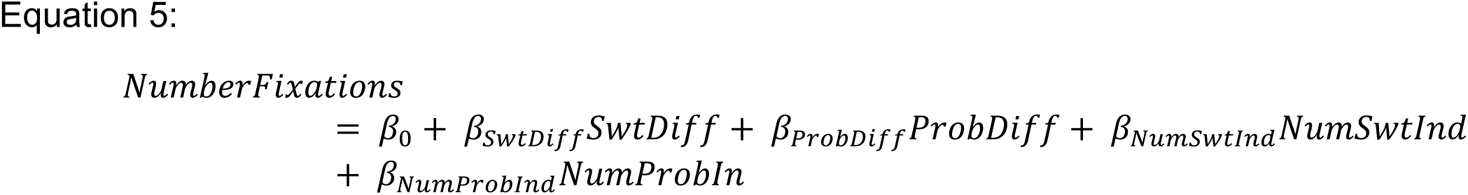

where SwtDiff represents the difference between the sweetnesses of the two options (0-4 a.u.), ProbDiff represents the difference between the probabilities of the two options (0-4 a.u.), NumSwtInd represents the number of sweetness bars that were indirect (0–2), and NumProbInd represents the number of probability bars that were indirect (0–2). All predictors were mean centered (ranges above are pre-mean-centering).

##### Regression on fixation duration

To quantify the effects of task variables on the duration of fixations, we used a linear regression model as follows:

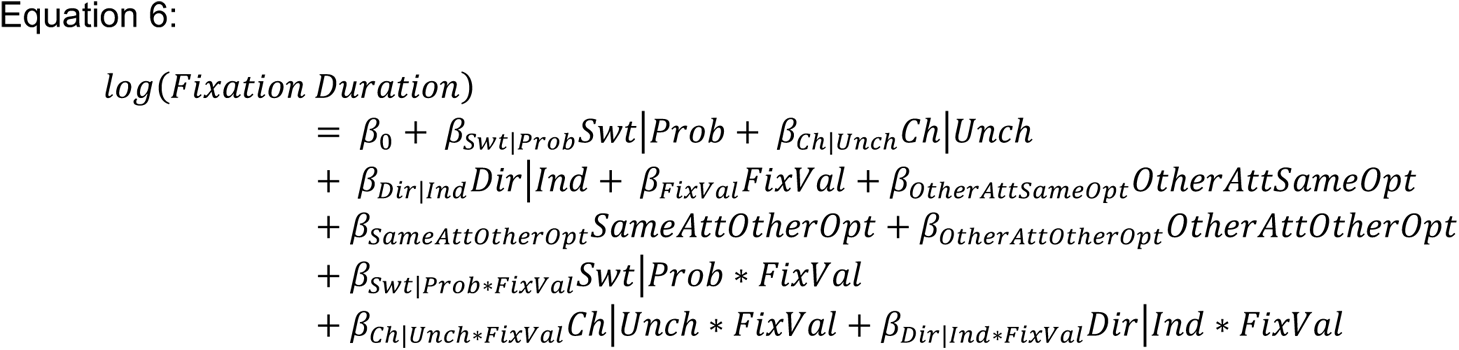

where Swt|Prob represents whether the fixated bar was for sweetness or probability (1, −1), Ch|Unch represents whether the fixated bar was part of the chosen or unchosen option (1,-1), Dir|Ind represents whether the fixated bar was the direct or indirect mapping (1,-1), and FixVal represents the attribute value of the fixated bar (1-5). OtherAttSameOpt represents the ordinal magnitude of the other attribute of the same option as the fixated bar (1-5), SameAttOtherOpt represents the ordinal magnitude of the same attribute as the fixated bar in the other option (1-5), and OtherAttOtherOpt represents the ordinal magnitude of the other attribute of the other option as the fixated bar (1-5). All predictors were mean-centered (ranges above are pre-mean-centering).

##### Hypothesis test on regression coefficients for gaze transitions

Value difference was regressed against the number of gaze transitions in two simple regressions. The first predicted the number of within-attribute gaze transitions within a trial (e.g., SwtA<->SwtB), and the second predicted the number of within-option gaze transitions within a trial (e.g., SwtA<->ProbA). The t-statistic for the two-sample hypothesis test that the slopes of these regressions differed was calculated as

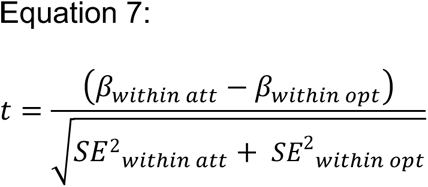

where SE represents the standard error of the regression coefficient (67).

## Acknowledgments

We would like to thank Mark Baxter, Peter Rudebeck, and Feng-Kuei Chiang for their comments on the manuscript. The research was supported by R01MH134845 and the Pew Biomedical Scholars Program to ELR and F31MH127901 to AQP.

---

**Supplemental Figure 1.**
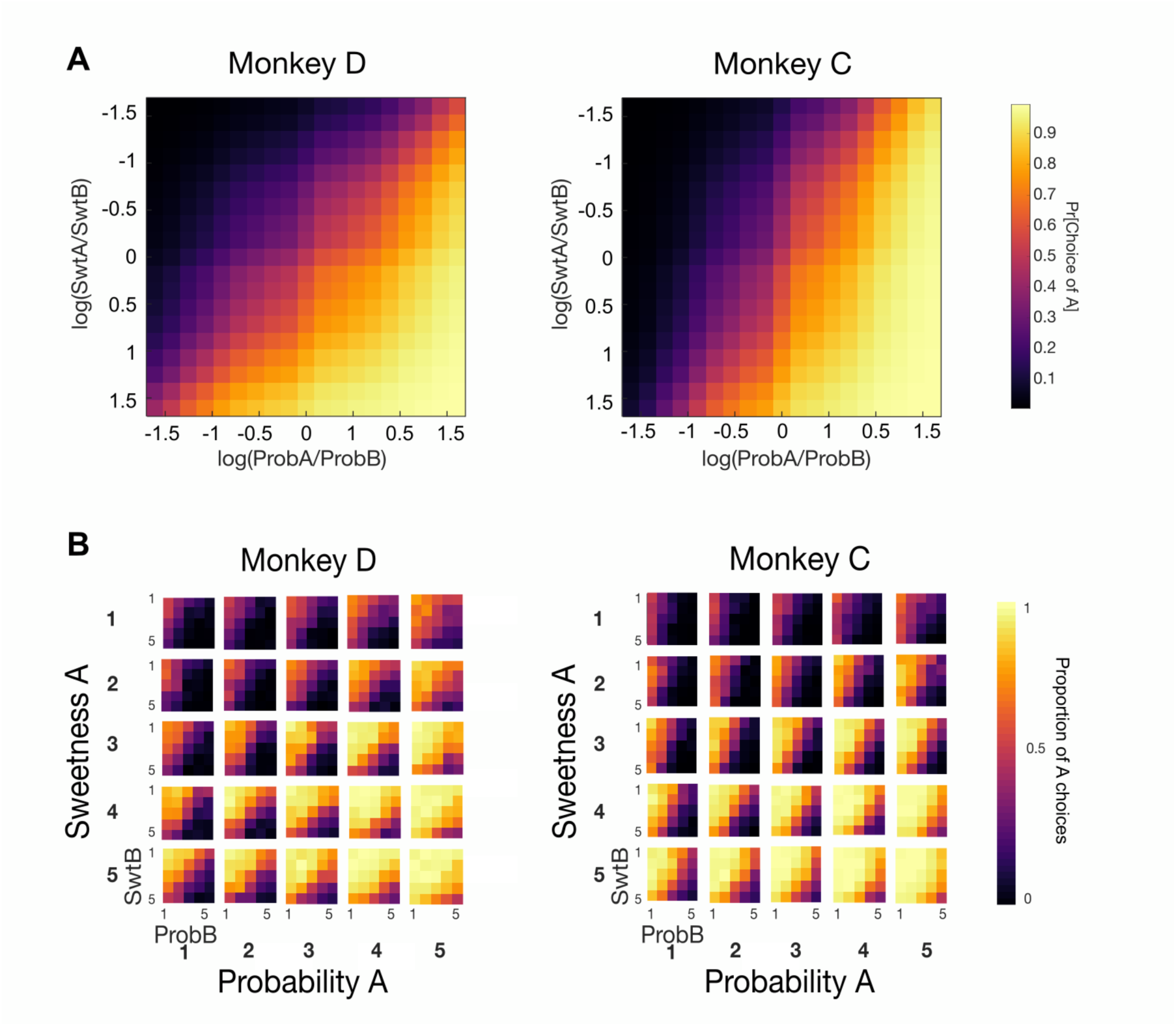
Monkeys use multiple attributes to make choices. **A**. Colormap of predicted choice probabilities from logistic regression on the log ratio of Sweetness A / B and Probability A/B (as in Fig 2D, but flattened) (Eqn. 3). Brighter colors represent greater frequency with which arbitrary option A was selected. N= 57082(D), 65384 (C). **B**. Colormaps of real choice frequencies. Each subplot shows choices when option A has sweetness magnitude 1 to 5 (subplots from top to bottom, y-axis) and probability magnitude of 1 to 5 (subplots left to right, x-axis). Within each subplot, magnitudes of option B vary in the same range (1-5 for each). SwtB/ProbB = sweetness/probability of option B.

**Supplemental Figure 2.**
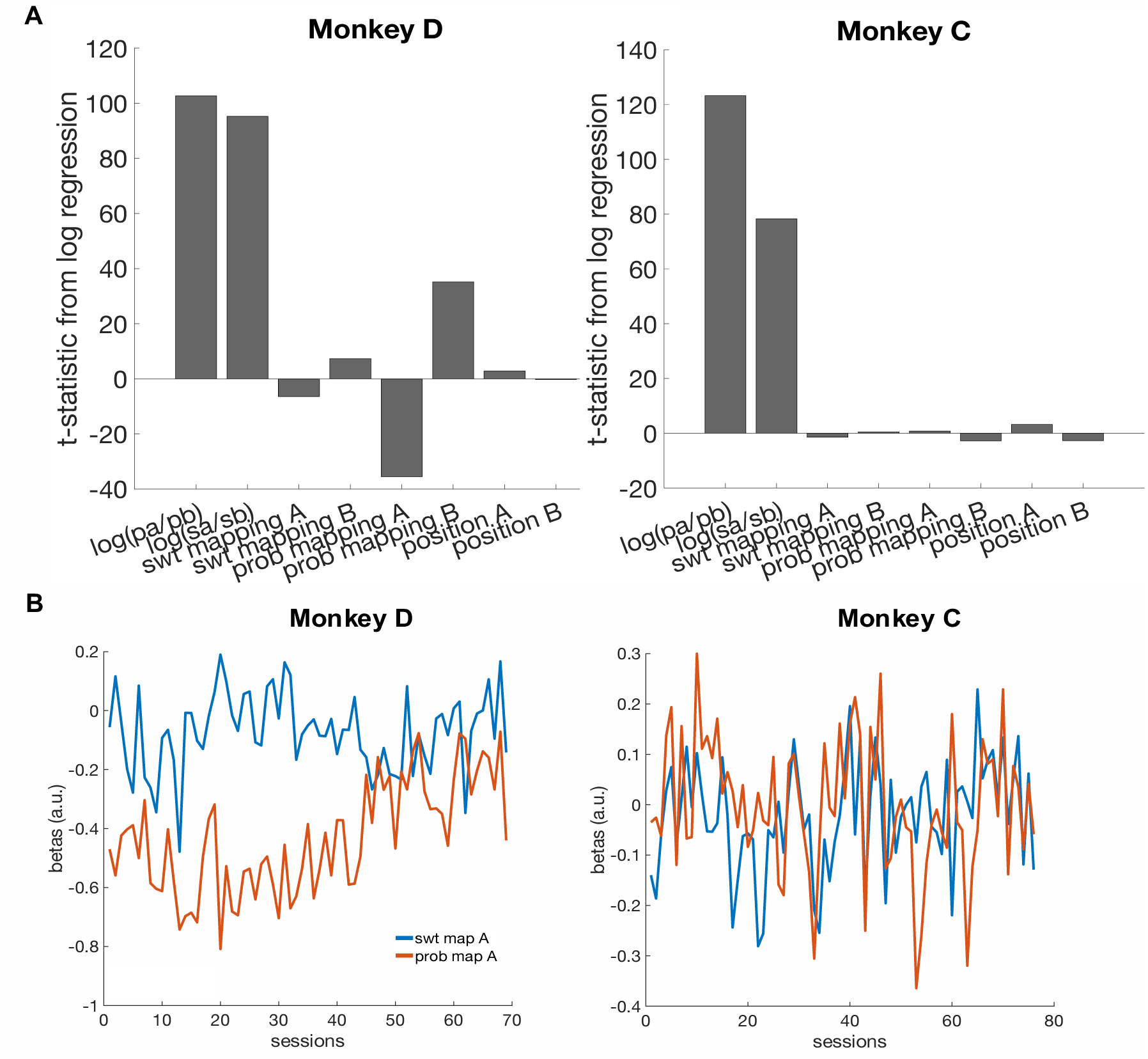
Regression coefficients from logistic regressions on choice. **A**. Regressions performed on concatenated data (Eqn. 1) found that the strongest predictors of choice are the relative magnitude of sweetness and probability of the two options. N=67082(D), 65384(C). **B.** Plot of regression coefficients for Sweetness / Probability mapping from the choice regression over sessions. Only mapping betas for option A are shown, as betas for A and B are roughly inverses of each other. Across sessions, the relative weighting of direct versus indirect mapping varied considerably. Monkey D initially preferred indirect Probability mappings, but this preference disappeared by the end of testing. Monkey C showed no consistent preference for direct or indirect mappings across testing.

**Supplemental Figure 3.**
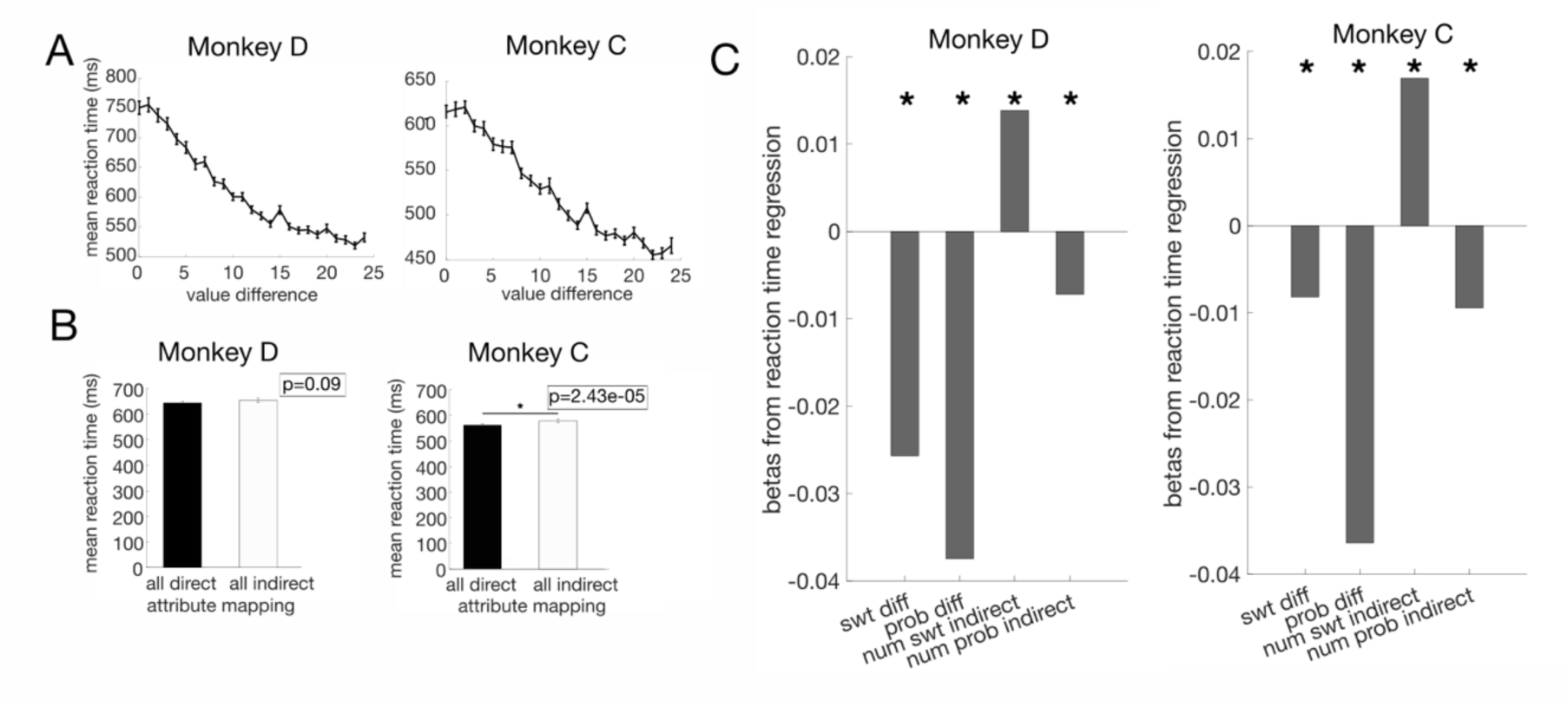
Reaction times (RTs) in multiple attributes choices. RTs were defined as the time between the appearance of choice options on the screen and the time when the touch bar was released to select an option. Average RTs across sessions were 672.20 ± 30.5ms for Monkey D, and 570.31 ± 23.2ms (95% CI) for Monkey C. RTs were shorter on trials with an objectively better option, compared to other trials, consistent with the idea that these are easier choices (Monkey D: 609.93 ± 11.68 ms (95% CI), Monkey C: 528.27 ± 10.33 ms (95% CI); two sample t-tests Monkey D: *p*=7.93×10^-27^; Monkey C: *p*=8.29×10^-40^). **A**. Average RTs across sessions varied with the difference in the expected value of the two options, defined as the ordinal sweetness x ordinal probability. Error bars = SEM. N= 69 (D) and 76 (C) (sessions). **B**. Reaction times were only weakly sensitive to direct/indirect attribute mapping in Monkey C (two-sample t-test). Error bars = SEM. N= 69 (D) and 76 (C) (sessions). **C**. Regression coefficients from a linear regression on reaction time. SwtDiff and ProbDiff are the differences between the ordinal value of each attribute (0-4). NumSwtInd and NumProbInd are the number of indirect mappings of each attribute in the trial (0-2). * p<0.01. N= 56889(D), 65322 (C).

**Supplemental Figure 4.**
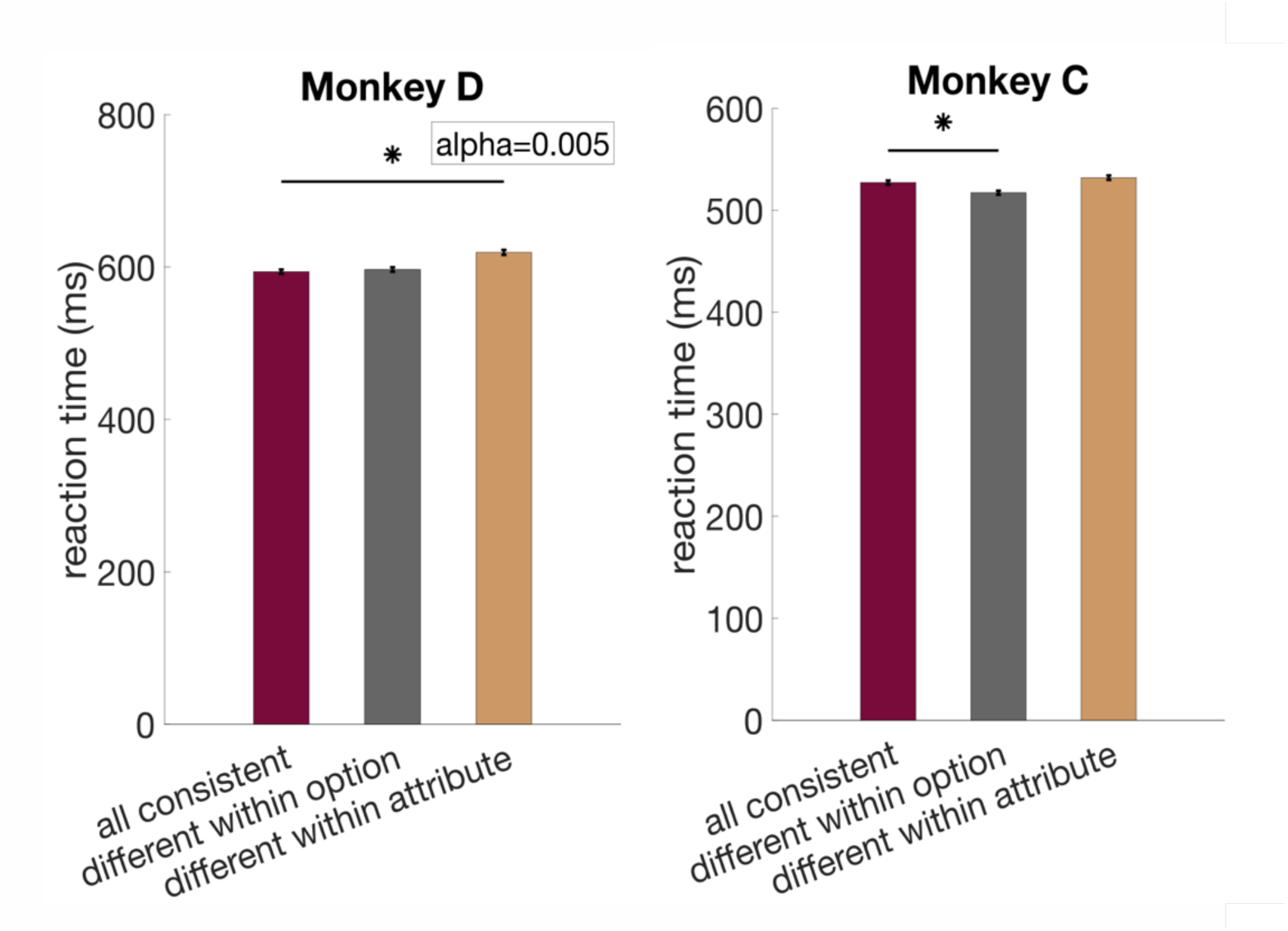
Reaction times split by attribute arrangement (as in Fig. 2E). Error bars = SEM. Statistics were performed on log-transformed reaction times. One-way ANOVA: Monkey D: F_2,10876_=22.07, p= 2.71×10^-10; Monkey C: F_2,12586_ =14.11, p=7.56×10^-7. Post-hoc comparisons, α=0.005; Monkey D: all consistent vs different within option: p= 0.98, all consistent vs different within attribute: p=4.77×10^-9; Monkey C: all consistent vs. different within option, p= 2.92×10^-4, all consistent vs different within attribute, p= 0.11. Due to the sample size, some comparisons reached significance, but the effect size is very small and may reflect spurious effects.

**Supplemental Figure 5.**
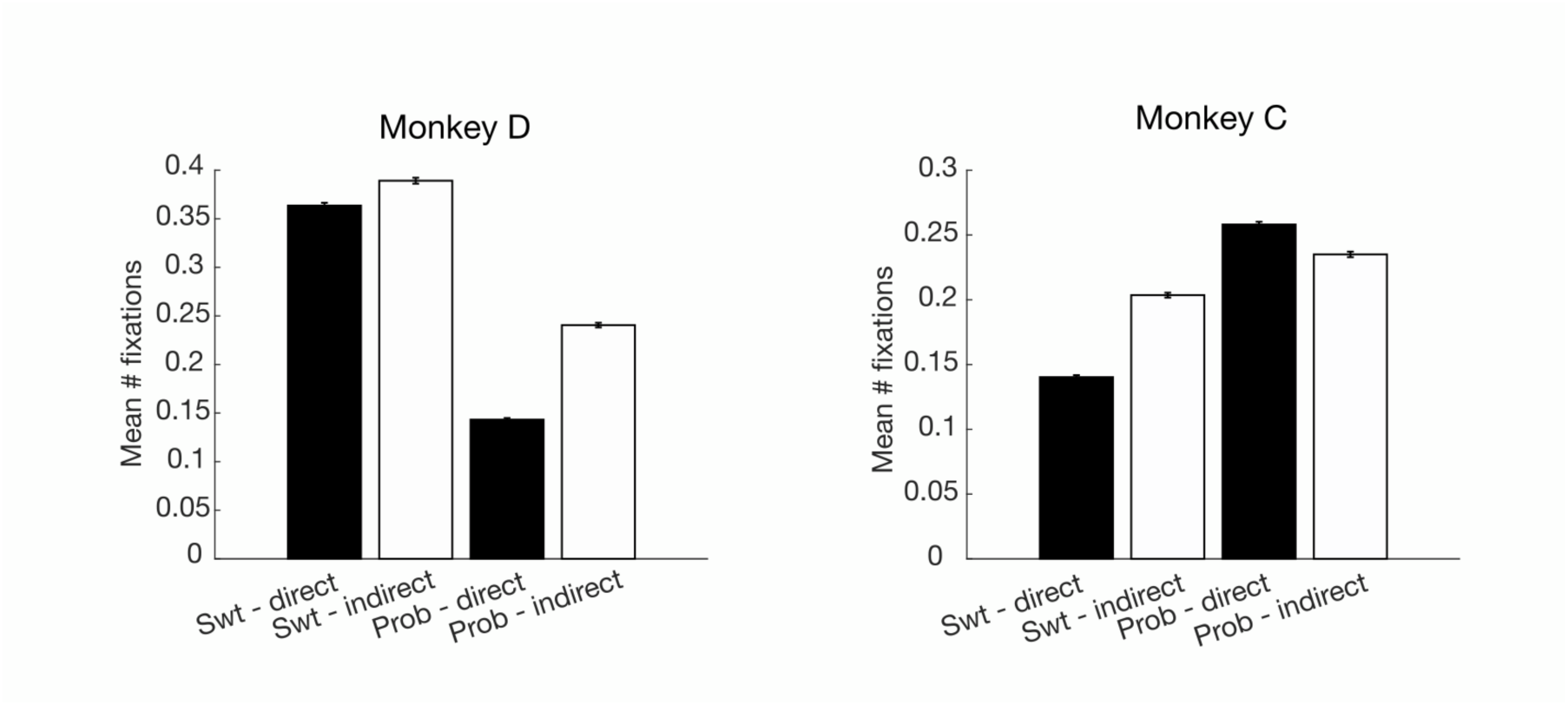
Mean number of fixations on attribute bars within a trial, split by direct/indirect mapping of each attribute. Fixations were counted between cue onset and choice. Error bars = SEM. N=51115 (D), 53727 (C) (trials).

**Supplemental Figure 6.**
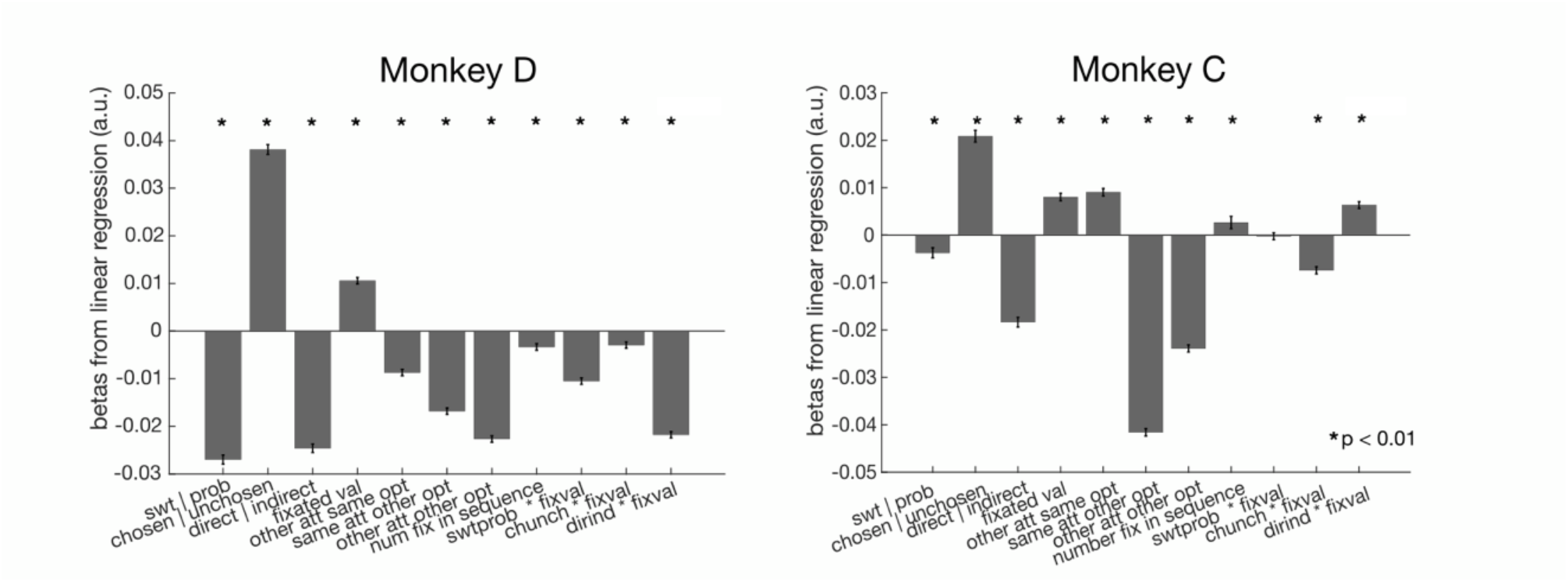
Regression coefficients from linear regressions on fixation duration (log(ms)). * = p<0.01. Error bars show standard error of the coefficients. N= 116043 (D), 89836 (C) (fixations).

